# The polarity specific nature of single session high-definition transcranial direct current stimulation to the cerebellum and prefrontal cortex on motor and non-motor task performance

**DOI:** 10.1101/2020.11.17.387217

**Authors:** Ted Maldonado, Jessica A. Bernard

## Abstract

The cerebellum has an increasingly recognized role in higher order cognition. Advancements in noninvasive neuromodulation techniques allows one to focally create functional alterations in the cerebellum to investigate its role in cognitive functions. To this point, work in this area has been mixed, in part due to varying methodologies for stimulation, and it is unclear whether or not transcranial direct current stimulation (tDCS) effects on the cerebellum are task or load dependent. Here, we employed a between-subjects design using a high definition tDCS system to apply anodal, cathodal, or sham stimulation to the cerebellum or prefrontal cortex (PFC) to examine the role the cerebellum plays in verbal working memory, inhibition, motor learning, and balance performance, and how this interaction might interact with the cortex (i.e. PFC). We predicted performance decrements following anodal stimulation and performance increases following cathodal stimulation, compared to sham. Broadly, our work provides evidence for cerebellar contributions to cognitive processing, particularly in verbal working memory and sequence learning. Additionally, we found the effect of stimulation might be load specific, particularly when applied to the cerebellum. Critically, anodal simulation negatively impacted performance during effortful processing, but was helpful during less effortful processing. Cathodal stimulation hindered task performance, regardless of simulation region. The current results suggest an effect of stimulation on cognition, perhaps suggesting that the cerebellum is more critical when processing is less effortful but becomes less involved under higher load when processing is more prefrontally-dependent.

## Introduction

Historically, the cerebellum was thought to be primarily involved in motor function (Holmes, 1939) and motor learning (Ballard et al., 2019; Bernard & Seidler, 2013). However, work over the last several decades has also implicated the cerebellum in non-motor cognitive processing (Buckner, 2013; Desmond et al., 1997; E et al., 2014; Leiner et al., 1989, 1991; Rapoport et al., 2000; Schmahmann, 2018; Schmahmann & Sherman, 1998; Stoodley, 2012). Briefly, imaging work demonstrates a segregated functional topography in the cerebellum (King et al., 2019; Stoodley et al., 2012; Stoodley & Schmahmann, 2010) such that anterior lobules and lobules VIIIa and VIIIb are implicated in motor functioning and the posterior lobules are primarily associated with cognitive functioning (e.g., Stoodley, 2012; Stoodley et al., 2012a; Stoodley, Valera, & Schmahmann, 2012b; Stoodley & Schmahmann, 2009; King et al., 2019). The segregated functional topography is thought to be driven by closed-loop cerebello-thalamo-cortical circuits (Bernard, Orr, & Mittal, 2016; Dum & Strick, 2003; Kelly & Strick, 2000; Palesi et al., 2015; Ramnani, 2006; Sen, Kawaguchi, Truong, Lewis, & Huang, 2010; Stoodley, Valera, & Schmahmann, 2012b; Stoodley & Schmahmann, 2009).

Though the literature implicating the cerebellum in non-motor cognitive processing is growing, little work examines the relative necessity of the cerebellum in executing cognitive tasks. Most work in this regard is limited to findings using lesion patients (Schmahmann, 1991; Schmahmann & Sherman, 1998; Timmann et al., 2010; Timmann & Daum, 2007). However, advances in noninvasive brain stimulation techniques, such as transcranial direct current stimulation (tDCS), allow us to create temporary lesions that further our understanding of cerebellar function (Ferrucci et al., 2015). tDCS sends either a positive (anodal) or negative (cathodal) current through the scalp with one electrode pad and a second pad is used to receive the current. Anodal stimulation is thought to increase cortical excitability, and cathodal stimulation decreases excitability (Brunoni et al., 2012; Nitsche & Paulus, 2000; Priori et al., 1998) of the underlying cortical area. Cerebellar tDCS has the potential to be particularly informative, as one can increase or decrease the cerebellum’s influence on cognitive processes, providing insight into the relative necessity of the cerebellum during task performance.

Critically, evidence suggests that cerebellar tDCS modulates task performance across multiple cognitive domains (Ferrucci & Priori, 2014; Oldrati & Schutter, 2018), though motor learning has been the primary focus of research with this technique (for a review see Buch et al., 2017). Recent work examining motor learning found polarity specific effects of stimulation over the lateral posterior cerebellum, such that cathodal stimulation improved initial sequence learning, and anodal stimulation hindered performance, relative to sham (Ballard et al., 2019). Similar findings were found in an ankle adaptation task (Shah et al., 2013). However, other studies have found the opposite polarity effects during motor learning paradigms (Block & Celnik, 2013; Cantarero et al., 2015; Galea et al., 2011; Hardwick & Celnik, 2014).

To date, tDCS work examining non-motor cognitive processes is limited, mixed and primarily focused on verbal working memory. Cathodal stimulation to the cerebellum has improved task performance on verbal working memory (Pope & Miall, 2012) and inhibition tasks (Mannarelli et al., 2019; Wynn et al., 2019), but has also resulted in performance decrements (Boehringer, Macher, Dukart, Villringer, & Pleger, 2013; Ferrucci et al., 2008) and increased response variability (Spielmann et al., 2017). Anodal cerebellar stimulation has been shown to decrease task performance for verbal working memory tasks (Ferrucci et al., 2008). It should also be noted that there is a growing literature demonstrating no effects of stimulation on cognitive task domains including implicit learning (Steiner et al., 2016; Verhage et al., 2017), working memory (Maldonado et al., 2019; van Wessel et al., 2016), probabilistic classification learning (Majidi et al., 2017), and inhibition (Maldonado et al., 2019).

Together, studies using cerebellar tDCS have been inconclusive with respect to the role of the cerebellum in cognitive processing. There are two notable possibilities for why this might be. First, much of the cerebellar tDCS work uses either a cathodal or anodal stimulation condition and one sham stimulation condition over the cerebellum. It is possible that much of the findings are mixed because there is an underlying polarity specificity by task interaction. Optogenetic work suggests polarity specific change in cerebellar activity following stimulation (Grimaldi et al., 2016), but little work has examined whether this polarity specificity is consistent across task types (i.e motor versus non-motor, inhibition versus working memory, etc.). It is possible that anodal stimulation improves task performance for some tasks but hinders performance in others, even within a broader task domain, as different component processes might be required to complete the task. For instance, does stimulation affect inhibitory processes differently when updating continually for a n-back task versus in chunks during a Sternberg task? But, to date, there are few direct comparisons of these active stimulation conditions on cognitive tasks.

Second, much of the non-motor cerebellar work ignores how performance changes in relation to the PFC, though there are known connections between the two brain regions (Bernard et al., 2016; Diedrichsen et al., 2019; King et al., 2019; Palesi et al., 2015; Sen et al., 2010). Regional interactions between the cerebellum and PFC on cognitive processing might be crucial in understanding how performance might change, as past work suggests the cerebellum might only be needed during certain aspects of cognitive processing, as a mechanism of support for frontally-driven processes (Filip et al., 2019). That is, there may be cortical compensatory processes at play that is resulting in null findings. Further, animal models suggest the cerebellum might be more involved in automatic processing (Ramnani, 2014), when a specific procedure is well learned. Therefore, cerebellar tDCS might reveal these specific interactions between the cerebellum and PFC, highlighting the relative necessity of the cerebellum during non-motor cognitive processing.

Therefore, we implemented a between-subjects study that used cathodal, anodal and sham stimulation, which was applied using high definition (HD) tDCS, an advancement in non-invasive neuromodulation that is thought to be more precise in targeting specific brain regions. This precision allows one to investigate both the cerebellum and PFC across both motor and cognitive tasks to provide insight into the necessity of the cerebellum for cognitive processing and how this necessity might differ from motor processing. Thus, we examined the role of the cerebellum in cognition and if there are polarity and region-specific outcomes in task performance, when compared to the PFC. Broadly, we predict performance increases following cathodal stimulation and performance decreases after anodal stimulation to the cerebellum. Though the literature is mixed, recent optogenetic work in mice has suggested directionality when predicting how stimulation will affect task performance. Grimaldi et al., (2016) suggests that anodal stimulation excites the inhibitory Purkinjie circuit which results in an increase in inhibitory function on the deep cerebellar nuclei (DCN) resulting in a decrease in cerebellar output, ultimately decreasing task performance. This is observed in behavioral work in humans as well (Ballard et al., 2019; Pope & Miall, 2012). Alternatively, it is believed that cathodal stimulation inhibits the inhibitory Purkinjie circuit which results in a decrease in inhibitory function on the DCN, resulting in an increase in cerebellar output, and ultimately increasing task performance. This finding will be used as a baseline for our predictions.

As such, for cognitive tasks, we predict increases in accuracy and decreases in reaction time following cathodal stimulation to the cerebellum. Further, we predict decreases in accuracy and increases in reaction time following anodal stimulation to the cerebellum. We also predict that these outcomes will be evident during low load trials in the Sternberg task, and during congruent trials in the Stroop task, particularly after cerebellar stimulation, as the cerebellum is thought to be involved in more automatic processing (Doyon, Gabitov, Vahdat, Lungu, & Boutin, 2018; Karni et al., 1998; Ramnani, 2014). Furthermore, we expect the same effects when stimulation is applied to the PFC; however, we predict these outcomes will be more evident during higher load trials in the Sternberg task, and during incongruent trials in the Stroop task, as the PFC is thought to be more involved in processing where cognitive demand is high (Braver et al., 1997; Rypma & D’Esposito, 1999). We predict similar outcomes when examining sequence learning data, in light of our recent findings (Ballard et al., 2019). That is, cathodal stimulation to the cerebellum will improve learning, particularly in early learning phases, and anodal stimulation to the cerebellum will hinder early learning.

In addition to the tasks mentioned above, we also had participants complete a balance task as the cerebellum is key in making postural adjustments to maintain balance (Ito, 2006). Critically, past work has indicated postural sway as an indicator of cerebellar function (Bernard et al., 2014) and is used here to gain a better insight into the effect of tDCS on cerebellar function on a motor specific task. We will use changes in area, variability and complexity of the center of pressure (COP) to determine changes in cerebellar function (Oliveira et al., 1996). Therefore, we predict cathodal stimulation to the cerebellum will reduce sway area and variability in sway, whereas anodal will increase sway area and variability in a balance task that will quantify postural sway. Regarding PFC stimulation, we predict similar outcomes, such that cathodal stimulation will improve performance when tasks demands are high, and anodal stimulation will create performance decrements.

To examine these hypotheses, participants were placed in one of three stimulation conditions (anodal, cathodal, or sham) and completed both motor (balance, sequence learning) and non-motor (Sternberg, stroop) tasks to better understand the necessity of the cerebellum plays in motor function and cognition. Further, stimulation was applied to either the cerebellum or the PFC, to understand regional differences in the impact of tDCS on cognitive processing, ultimately providing better insight into the role the cerebellum plays in cognition, in relation to the PFC.

## Methods

### Participants

One hundred and sixty-three right-handed undergraduate students from Texas A&M University participated in this study for partial course credit in an Introduction to Psychology course. Data from 7 participants were not used because of a computer error that resulted in incomplete data sets. Thus, 156 right-handed participants (105 female) ages 18 to 22 (*M*=18.86 years, *SD*=0.94) were included in the analysis. As this is a between-subjects analysis, there were 26 participants per region, per stimulation condition (6 total groups). All procedures completed by participants were approved by the Texas A&M University Institutional Review Board and conducted according to the principles expressed in the Declaration of Helsinki.

### Procedure

Data was collected in a single 90-minute visit. Participants completed a demographic survey and the Edinburgh Handedness Inventory (Oldfield, 1971) to confirm right-handedness, followed by tDCS (see below for details). After the tDCS administration, participants then completed a balance task, along with a computerized Stroop (Stroop, 1935), Sternberg (Sternberg, 1966), and sequence learning (Kwak et al., 2012) task, in a pre-determined random order (for more details, see below). Stimulation was administered in a single-blind manner, wherein participants were blinded to the stimulation type.

#### tDCS Stimulation Parameters

Participants were first fitted with a cap that followed the 10-20 electrode system. The cap was centered over Cz. We used the Soterix MxN HD-tDCS system, which allows for nine electrodes to be applied to the scalp, as such a montage can increase stimulation focality. Unlike traditional tDCS which relies on two pads for stimulation, high definition (HD) tDCS provides increased precision by using multiple electrodes (Datta et al., 2016; Huang et al., 2017), that will optimally target a specific region of interest. Electrode montages are determined using the accompanying Soterix Neurotargeting HD-Targets Software (Datta et al., 2016; Huang et al., 2017). In a typical multi-electrode montage, a ring is created around the central electrode(s) (Datta et al., 2009, 2012; Dmochowski et al., 2011; Villamar et al., 2013). The ring applies the opposite current to localize the current applied by the central electrode. In this study, two or three electrodes, depending on brain region, set the current (cathodal, anodal, or sham), and this current was contained by applying the opposite current using the remaining electrodes (See Table 1 and Figure 1). A ninth electrode was used as a return. Specially designed elastic straps were used to extend the cap beyond a 64-channel montage to ensure cerebellar stimulation. The specific electrode locations and current at each site are presented in Table 1. Figure 1 presents the modelled current flow targeting the right lateral posterior cerebellum and Figure 2 presents the modelled current flow targeting the left dorsal lateral PFC.

**Table 1.** Current intensities (in mA) and locations for cathodal stimulation to the cerebellum and PFC. Anodal stimulation used the same locations and intensities; however, the directions were flipped, meaning negative currents are positive during anodal stimulation and vice versa.

|  | Cerebellar Stimulation |  | PFC Stimulation |  |
| --- | --- | --- | --- | --- |
|  | Location | Current | Location | Current |
| **1** | P2 | 0.1135 | FP1 | 1.0062 |
| **2** | PO10 | -0.1551 | F3 | -1.757 |
| **3** | O10 | -0.3618 | F4 | 0.2221 |
| **4** | Ex3 | 0.1956 | P3 | 0.0761 |
| **5** | Ex4 | -1.4832 | C2 | 0.0183 |
| **6** | Ex5 | 0.4066 | CP2 | 0.1049 |
| **7** | Ex12 | 0.22 | F5 | -0.8243 |
| **8** | Ex14 | 0.7886 | TP7 | 0.0699 |
| **C** | Nk1 | 0.2757 | F9 | 0.5024 |

**Figure 1.**
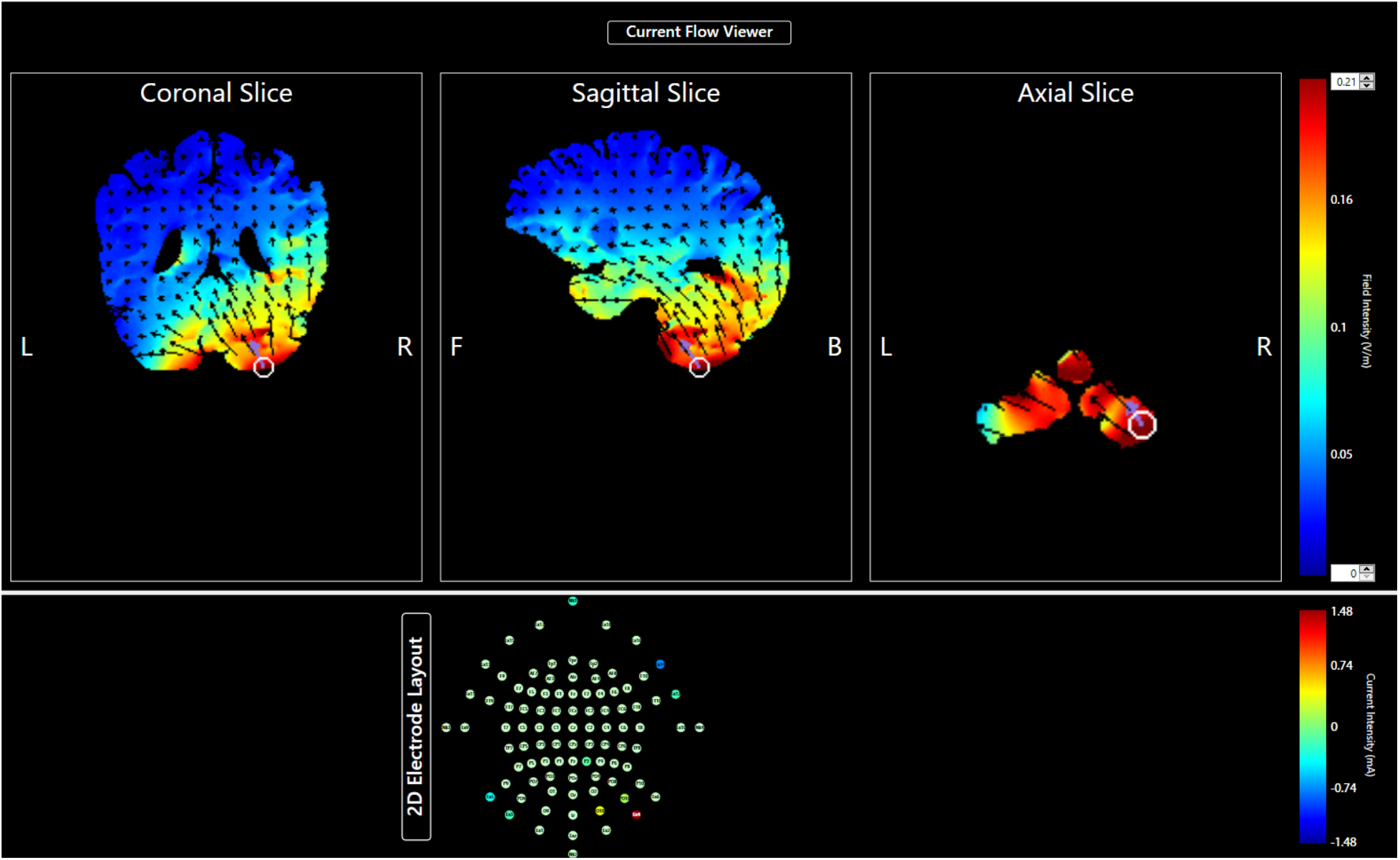
Modelled current flow and intensity (in V/m ranging from −1.48 to 1.48) montage using Soterix Targeting software to target the right cerebellum. Cooler colors indicate lower intensities while warmer colors indicate higher intensities. *Note*. L=Left, R=Right, F=Front, B=Back.

**Figure 2.**
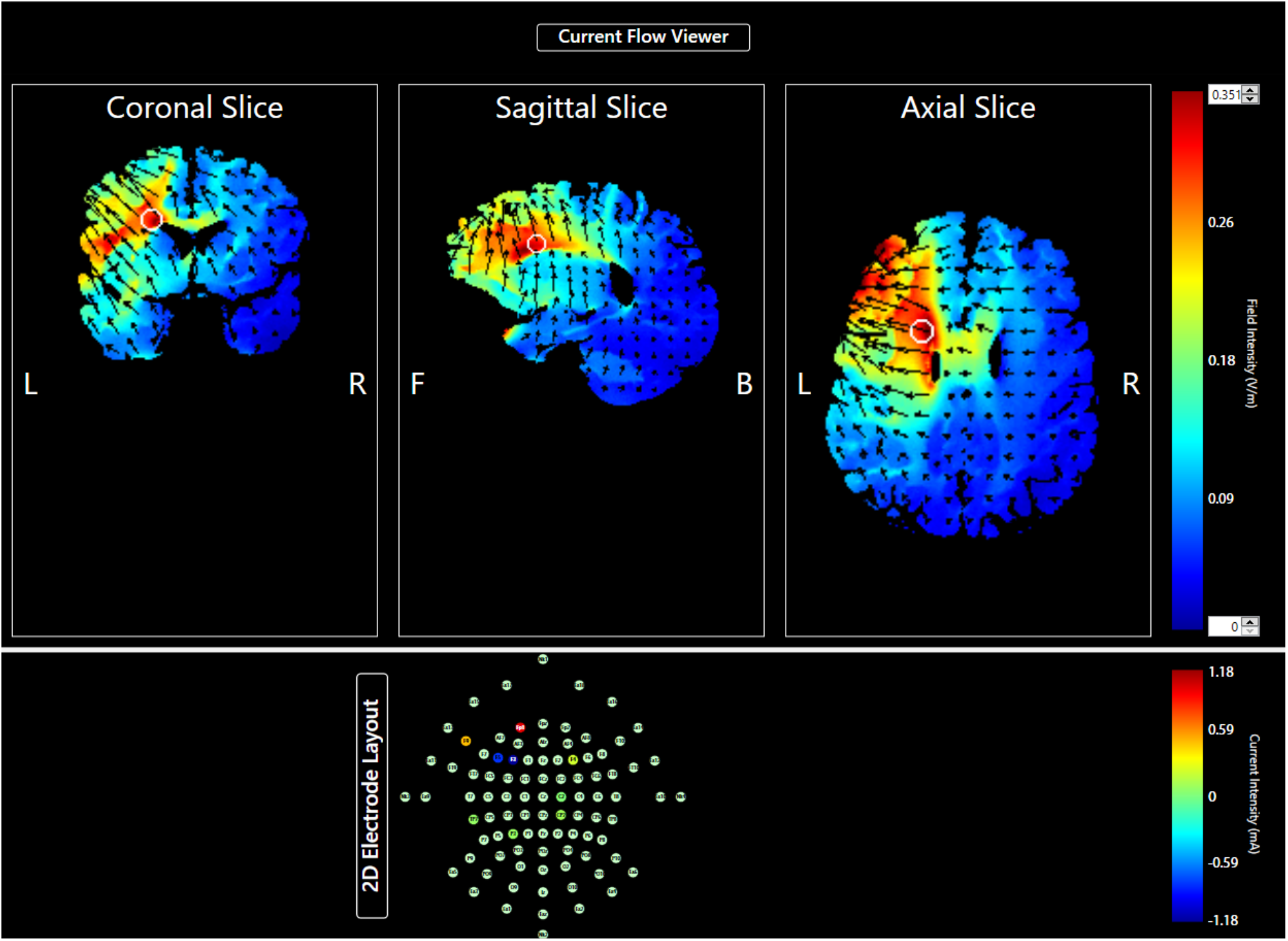
Modelled current flow and intensity (in V/m ranging from −1.48 to 1.48) montage using Soterix Targeting software to target the left PFC. Cooler colors indicate lower intensities while warmer colors indicate higher intensities. *Note*. L=Left, R=Right, F=Front, B=Back.

Following capping, stimulation was set to 0.1 mA for one minute to ensure the electrodes made a good connection with the scalp and that the appropriate impedance levels for stimulation (less than 100 KOhm) were reached. Adjustments, such as adding more gel to improve the bridge between the scalp and electrode, were made for electrodes with an impedance over 100 KOhm and the impedance was rechecked. Once impedance levels were at appropriate levels, participants completed a 20-minute stimulation session at 2 mA (Ferrucci et al., 2015; Grimaldi et al., 2014, 2016), using the currents presented in Table 1. During the stimulation conditions, the currents gradually increased for 30 seconds, maintained intensity for 20 minutes, and gradually decreased for 30 seconds. During sham conditions, the currents gradually increased for 30 seconds until currents were reached, then gradually decreased for 30 seconds directly before and after the 20-minute session. There was no additional stimulation during the 20-minute session. Stimulation was followed by the completion of the cognitive and motor tasks.

#### Behavioral Paradigms

The order of the following tasks was counterbalanced across participants and conditions to mitigate the impact of time after stimulation on our effects. In total, the tasks took approximately 50 minutes to complete, including time for instructions.

##### Stroop

The Stroop task was administered via computer using a preprogrammed script from Experiment Factory (Sochat, 2018). Participants were shown a series of color words (RED, BLUE, GREEN) and told to identify the ink color via button press. The ink color and the word were congruent in 50% of trials, such that the word ‘BLUE was written in blue ink. The remaining trials were incongruent, such that the word ‘RED” might be written in green ink. Stimuli were shown for 1500 milliseconds and were instructed to respond quickly and accurately. Participants completed 120 trials. Dependent variables were average reaction time for all trials, accuracy, and the Stroop effect. The Stroop effect refers to the interference experienced when naming the ink color during incongruent trials.

##### Sternberg

The Sternberg Task was administered via computer using Presentation Software (Version 18.0, Neurobehavioral Systems, Inc., Berkeley, CA, www.neurobs.com). At the beginning of each block, a participant was shown and told to remember a string of either one, three or five capitalized letters for five seconds, representing low, medium, and high loads, respectively. Following the presentation of the study letters, participants were shown individual lower-case letters at a rate of one letter per second. Participants were told to indicate whether the letter was one of the original letters shown at the beginning of the block. Participants completed two blocks for each load level for a total of six blocks. Each block had 25 trials, for a total of 150 trials. Dependent variables were average reaction time for correct trials and accuracy.

##### Sequence

The sequence task was administered via computer using Presentation Software (Version 18.0, Neurobehavioral Systems, Inc., Berkeley, CA, www.neurobs.com) and was based on a paradigm used by Kwak and colleagues (2012). Participants were shown four empty rectangles and instructed to indicate the location of the rectangle that was filled in via button press. Each stimulus was shown for 200 ms and the participant had 800 ms to respond, before the next stimulus appeared. Random blocks (R) had 18 trials and sequence (S) blocks had 36 trials. During sequence trials, a six-element sequence (2-4-1-3-2-4) was repeated six times for the participant to learn. The order of the task was as follows: R-S-S-S-R-R-S-S-S-R-R-S-S-S-R in which the first three sequence blocks are considered early learning, the central sequence block is middle learning, and the last sequence block is considered late learning. The sequence did not include any trills (e.g. 747) or repeats (e.g. 777). Dependent variables included mean reaction time for correct trials and average total accuracy to estimate learning.

##### Balance

To assess balance, as quantified by bodily sway, data was collected using an Advanced Mechanical Technology Incorporated (AMTI) Accusway (Watertown, MA) force platform. Participants were instructed to remove their shoes and stand on the force plate for two minutes per condition, with a sample rate of 200 samples per second. Participant completed four conditions. These include eyes open, open base (EOOB); eyes open, closed base (EOCB); eyes closed, open base (ECOB); eyes closed, closed base (ECCB). During conditions where eyes were open, participants were instructed to focus on a cross on the wall in front of them. For each condition we recorded center of pressure (COP). COP and the 95% confidence interval of COP area were measured using principle component analysis (Bernard et al., 2014; Oliveira et al., 1996; Osborne et al., 2017). Additionally, a 9th order Butterworth filter with a 20 Hz cutoff frequency was applied to isolate low-frequency sway. There were five dependent variables. The mean area of sway, the standard deviation in sway in the X and Y direction, and the complexity of the sway pattern in the X and Y direction were computed using Matlab (Kent et al., 2012).

### Data Processing and Analysis

All statistical analyses were completed in R (Team, 2018), using the “lme4” (Bates et al., 2015) package, with p-values estimates determined using the “lmerTest” package (Kuznetsova et al., 2017).

Stroop, Sternberg, sequence, and balance task data were analyzed using liner mixed effects models using restricted maximum likelihood, as it produces unbiased estimates of variance and covariance parameters, ideal for mixed effect models with small samples. Several fixed factors were used across tasks, which include stimulation type (cathodal, anodal, or sham stimulation) and brain region (cerebellum, PFC). Subject was used as a random effect for all models. For Sternberg, load (low, medium, and high) was included as a fixed factor. Congruency (congruent and incongruent) was included as a fixed factor for the Stroop task. Phase (early, middle, late) was included as a fixed factor for the sequence task, with all random trials included for comparison. A model was completed for each dependent variable listed above. For balance, the standing condition (EOOB, EOCB, ECOB, ECCB) was included as a fixed factor. There were five dependent variables: Sway area, standard deviation in sway in the X (SDx) and Y (SDy) direction and the complexity of the sway pattern in the X (ALPHAx) and Y (ALPHAy) direction.

All results were evaluated with a statistical threshold wherein p<.05 was used as the cut-off for significance. When necessary, significant effects were followed up by comparisons of estimated marginal means with Bonferroni-corrected *p*-values using the emmeans package (Lenth et al., 2018).

## Results

### Stroop

Mean reaction times for the Stroop task can be found in Table 2. We first analyzed reaction time data and only found an effect of congruency [(*F*(1, 17,726) = 1,446.25, *p* < 0.001)], where reaction times were higher for incongruent trials than congruent trials (*p* < .0001). We also found a trend level effect of region [(*F*(1, 149) = 3.90, *p* = .050)], in which reaction time was quicker in the cerebellum group compared to the PFC group (*p =* .048). No other effects reached significance (*F*s<01.58, *p*s>.21).

**Table 2.** Mean RT and accuracy for the Stroop task by Congruency, Region, and Stimulation Condition.

| Congruency | Region | Stimulation | Reaction Time (ms) |  | Accuracy |  |
| --- | --- | --- | --- | --- | --- | --- |
|  |  |  | Mean | SD | Mean | SD |
| Congruent | Cerebellum | Anodal | 668.64 | 184.99 | 0.99 | 0.10 |
| Congruent | Cerebellum | Cathodal | 681.33 | 200.47 | 0.99 | 0.12 |
| Congruent | Cerebellum | Sham | 665.27 | 195.61 | 0.99 | 0.10 |
| Congruent | PFC | Anodal | 690.61 | 209.53 | 0.99 | 0.08 |
| Congruent | PFC | Cathodal | 701.86 | 200.01 | 0.98 | 0.13 |
| Congruent | PFC | Sham | 710.19 | 193.17 | 0.99 | 0.08 |
| Incongruent | Cerebellum | Anodal | 773.85 | 229.03 | 0.95 | 0.23 |
| Incongruent | Cerebellum | Cathodal | 787.76 | 232.12 | 0.94 | 0.24 |
| Incongruent | Cerebellum | Sham | 766.53 | 223.59 | 0.95 | 0.22 |
| Incongruent | PFC | Anodal | 804.47 | 239.03 | 0.94 | 0.24 |
| Incongruent | PFC | Cathodal | 812.24 | 234.63 | 0.94 | 0.24 |
| Incongruent | PFC | Sham | 819.46 | 232.36 | 0.94 | 0.23 |

Mean accuracy scores for the Stroop task can be found in Table 2. Similarly, when examining accuracy scores, we found an effect of congruency [(*F*(1,17,718.3) = 312.07, *p* < 0.001)], such that accuracy was higher for congruent trials than incongruent trials (*p* < .0001). No other effects reached significance (*F*s<1.29, *p*s>.28). Lastly, we found no significant effects of stimulation when examining the Stroop effect (*F*s<0.91, *p*s>.40).

### Sternberg

#### Reaction time

Mean reaction times for the Sternberg task can be found in Table 3 and are depicted visually in Figure 3. When examining the fixed effects of reaction time, there was a significant effect of load [(*F*(2, 16614.7) = 103.06, *p* < 0.001)], such that reaction times for each load condition were significantly different from each other (*p*s<.024). This demonstrates the increase in difficulty associated with increased load. Further, a significant stimulation by load interaction emerged [(*F*(4, 16614.7) = 7.05, *p* < 0.001)]. Specifically, reaction time under high load is significantly slower than medium load following anodal stimulation (*p* < 0.001), but not cathodal (*p* = 1.00) or sham stimulation (*p* = 1.00). Taken together, when collapsing across stimulation regions, anodal stimulation negatively impacted RT, particularly when cognitive processing is high.

**Table 3.** Mean RT and accuracy for the Sternberg task by Load, Region, and Stimulation Condition.

| Load | Region | Stimulation | Reaction Time (ms) |  | Accuracy |  |
| --- | --- | --- | --- | --- | --- | --- |
|  |  |  | Mean | SD | Mean | SD |
| Low | Cerebellum | Anodal | 648.19 | 131.08 | 0.93 | 0.26 |
| Low | Cerebellum | Cathodal | 658.15 | 143.89 | 0.90 | 0.30 |
| Low | Cerebellum | Sham | 632.80 | 139.18 | 0.91 | 0.29 |
| Low | PFC | Anodal | 638.40 | 137.56 | 0.91 | 0.28 |
| Low | PFC | Cathodal | 643.18 | 127.88 | 0.93 | 0.26 |
| Low | PFC | Sham | 657.09 | 141.79 | 0.92 | 0.28 |
| Medium | Cerebellum | Anodal | 676.69 | 138.58 | 0.85 | 0.35 |
| Medium | Cerebellum | Cathodal | 678.72 | 144.87 | 0.79 | 0.40 |
| Medium | Cerebellum | Sham | 661.96 | 151.37 | 0.82 | 0.38 |
| Medium | PFC | Anodal | 666.29 | 150.44 | 0.83 | 0.38 |
| Medium | PFC | Cathodal | 674.83 | 148.15 | 0.80 | 0.40 |
| Medium | PFC | Sham | 679.67 | 151.25 | 0.82 | 0.39 |
| High | Cerebellum | Anodal | 691.36 | 147.15 | 0.70 | 0.46 |
| High | Cerebellum | Cathodal | 681.16 | 167.51 | 0.69 | 0.46 |
| High | Cerebellum | Sham | 653.04 | 153.00 | 0.74 | 0.44 |
| High | PFC | Anodal | 696.37 | 156.09 | 0.74 | 0.44 |
| High | PFC | Cathodal | 670.01 | 161.12 | 0.72 | 0.45 |
| High | PFC | Sham | 685.71 | 156.28 | 0.75 | 0.43 |

**Figure 3.**
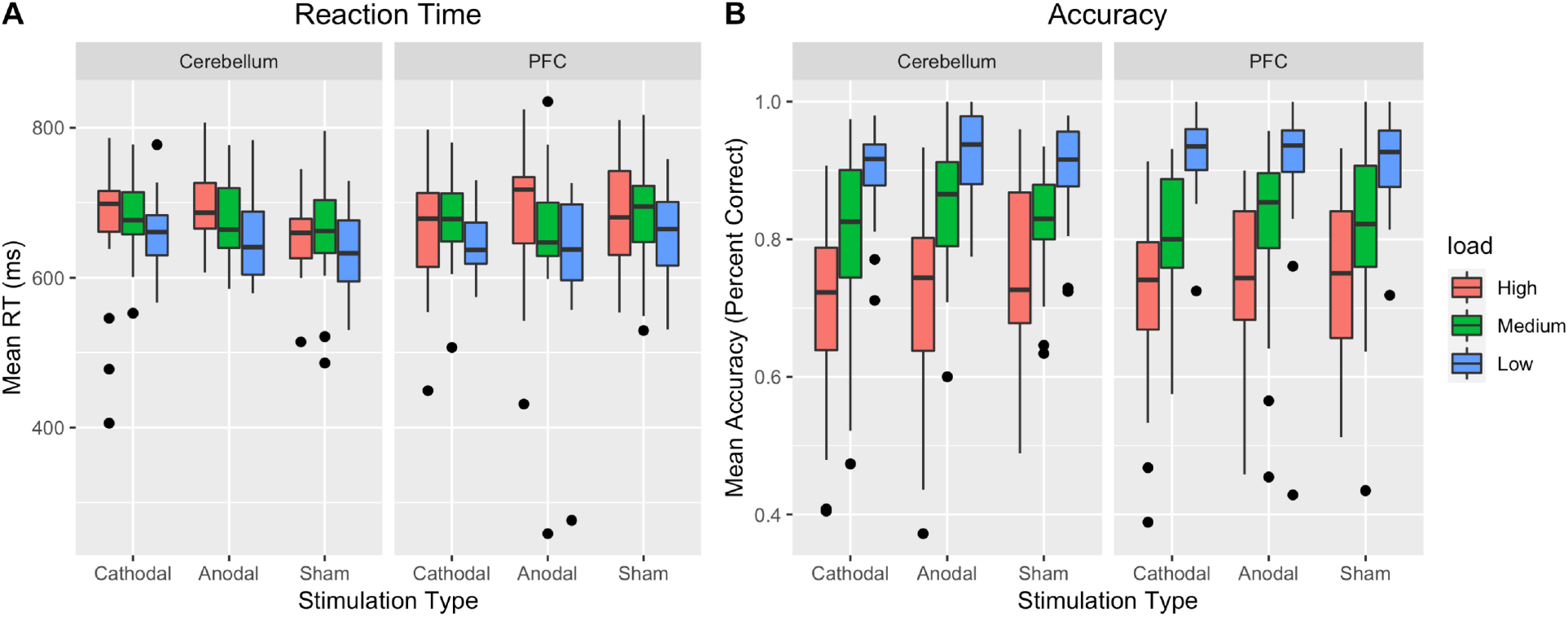
Mean RT and accuracy for the Sternberg task by Load, Region, and Stimulation Condition. Dots indicate outliers. Whiskers represent the interquartile range.

Further, a marginal load by region interaction emerged [(*F*(2, 16614.7) = 2.45, *p* = 0.087)]. We found that reaction time was significantly different across all load conditions in the PFC group (*p* < 0.01). In the cerebellar group, reaction time was significantly different between high and low (*p* < 0.01) and medium and low loads (*p* < 0.01). Critically, reaction times were not different, between high and medium load (*p* = 1.00). No other effects reached significance (*F*s<1.68, *p*s>.19).

#### Accuracy

We also examined the effects on accuracy (Table 3, Figure 3). There was an effect of load [(*F*(2, 20259) = 445.15, *p* < 0.001)], such that accuracy was best on low (*p* < 0.001) load, then medium load (*p* < 0.001), and then high load (*p* < 0.001). There was also a significant stimulation by load interaction [(*F*(4, 20259) = 3.97, *p* = .003)]. This interaction seems to be driven by marginal effects, such that accuracy was higher following anodal stimulation under medium load trials, compared to cathodal (*p* = .053). Additionally, accuracy was worse following cathodal stimulation under high load, compared to sham (*p* = 0.051).

There was also a marginally significant region by load interaction [(*F*(2, 20259) = 2.88, *p* =.056)], but follow up testing did not reveal any significant effects (*p*s > .138). No other effects reached significance (*F*s<1.68, *p*s>.15). Taken together, when collapsing across region, anodal stimulation improved accuracy under medium load, and cathodal stimulation hindered accuracy under high load.

### Sequence Learning

#### Reaction Time

Mean reaction times for the Sequence task can be found in Table 4 and depicted visually in Figure 4. First, we found a significant phase by region by stimulation interaction [(*F*(6, 53032) = 9.02, *p* < .001)]. Specifically, reaction time is slower for early (*p* = 0.003), middle (*p* = 0.016), and late (*p* < 0.001) learning, and random trials (*p* = 0.053), following cathodal stimulation to the cerebellum, when compared to sham stimulation (See Figure 4). Additionally, we found that reaction time is slower for late (*p* = .025) learning following cathodal stimulation to the cerebellum, when compared to anodal stimulation. No other effects were significant (*F*s<3.05, *p*s>.08).

**Table 4.** Mean RT and accuracy for the sequence learning task by Phase, Region, and Stimulation Condition.

| Phase | Region | Stimulation | Reaction Time (ms) |  | Accuracy |  |
| --- | --- | --- | --- | --- | --- | --- |
|  |  |  | Mean | SD | Mean | SD |
| Early | Cerebellum | Anodal | 350.46 | 114.47 | 0.94 | 0.24 |
| Early | Cerebellum | Cathodal | 385.50 | 126.83 | 0.93 | 0.26 |
| Early | Cerebellum | Sham | 330.47 | 104.01 | 0.93 | 0.26 |
| Early | PFC | Anodal | 332.87 | 100.54 | 0.94 | 0.24 |
| Early | PFC | Cathodal | 345.05 | 112.02 | 0.94 | 0.24 |
| Early | PFC | Sham | 346.45 | 108.00 | 0.94 | 0.24 |
| Middle | Cerebellum | Anodal | 346.01 | 132.28 | 0.92 | 0.28 |
| Middle | Cerebellum | Cathodal | 365.20 | 128.45 | 0.86 | 0.35 |
| Middle | Cerebellum | Sham | 315.55 | 106.03 | 0.90 | 0.30 |
| Middle | PFC | Anodal | 318.52 | 104.74 | 0.91 | 0.29 |
| Middle | PFC | Cathodal | 329.71 | 111.57 | 0.90 | 0.29 |
| Middle | PFC | Sham | 312.31 | 94.27 | 0.95 | 0.22 |
| Late | Cerebellum | Anodal | 328.71 | 133.03 | 0.86 | 0.35 |
| Late | Cerebellum | Cathodal | 370.34 | 132.25 | 0.85 | 0.36 |
| Late | Cerebellum | Sham | 297.37 | 93.22 | 0.88 | 0.32 |
| Late | PFC | Anodal | 312.07 | 108.64 | 0.90 | 0.30 |
| Late | PFC | Cathodal | 316.64 | 113.10 | 0.82 | 0.39 |
| Late | PFC | Sham | 305.07 | 97.69 | 0.94 | 0.24 |
| Random | Cerebellum | Anodal | 407.91 | 109.35 | 0.93 | 0.26 |
| Random | Cerebellum | Cathodal | 427.44 | 114.13 | 0.92 | 0.28 |
| Random | Cerebellum | Sham | 390.83 | 98.96 | 0.92 | 0.28 |
| Random | PFC | Anodal | 388.47 | 103.86 | 0.92 | 0.27 |
| Random | PFC | Cathodal | 400.63 | 108.04 | 0.91 | 0.29 |
| Random | PFC | Sham | 391.20 | 97.85 | 0.94 | 0.25 |

**Figure 4.**
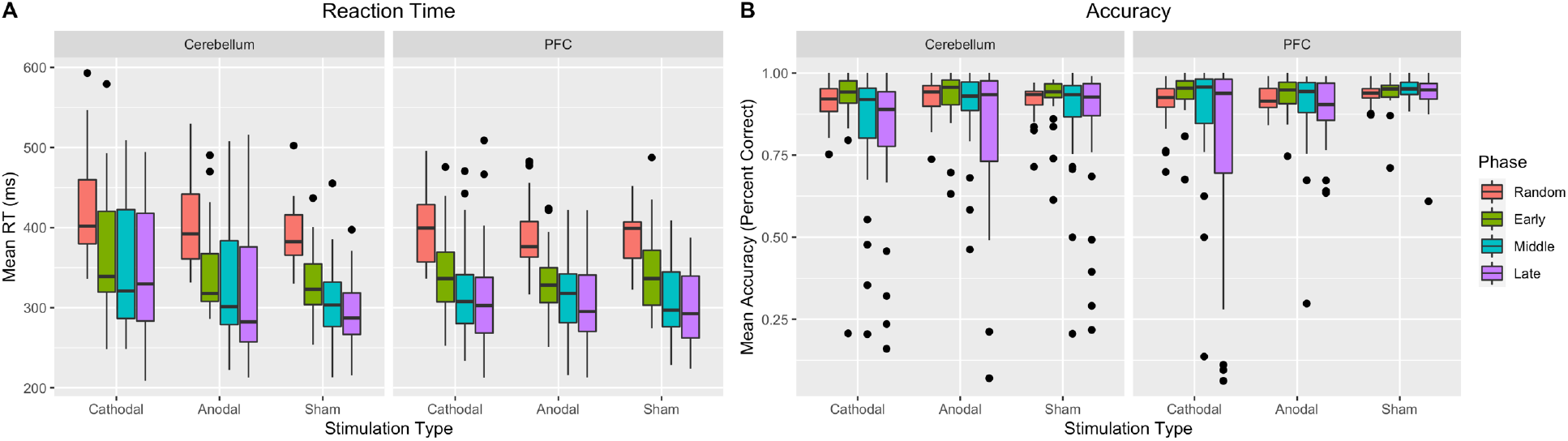
Mean RT and accuracy for the sequence learning task by Phase, Region, and Stimulation Condition. Dots indicate outliers. Whiskers represent the interquartile range.

Subsequent to the three way interactions, we found significant effect of learning phase [(*F*(3, 53,032) = 2,128.16, *p* < 0.001)], such that reaction times for early, middle, late, and random learning trials were all significantly different from one another (*p*s < .001). Additionally, we found a main effect of stimulation type [(*F*(2, 149) = 3.24, *p* = .042)], in which reaction times following cathodal stimulation were significantly slower than reaction times following sham stimulation (*p*=.040). Further, we did not find an effect of region [(*F*(1, 149) = 3.05, *p* = .083)].

We also found a significant phase by stimulation interaction [(*F*(6, 53032) = 5.49, *p* < 0.001)], in which reaction times were slower during middle (*p*=.033) and late (*p* = .009) learning phases, but not the early learning phase (*p* =.11), following cathodal stimulation, when compared to sham. Lastly, we unpacked the significant phase by region interaction [(*F*(3, 53,032) = 4.21, *p* = .006)]. This is driven by a longer reaction time in the middle (*p* = .037) and late (*p* =.053) phases of learning in the cerebellar stimulation group, relative to the PFC stimulation group. Taken together, cathodal stimulation to the cerebellum seemed to negatively impact reaction time across learning phases, whereas anodal stimulation does not seem to alter performance, regardless of stimulation location.

#### Accuracy

Mean accuracy scores for the Sequence task can be found in Table 4 and depicted visually in Figure 4. First, we found a phase by stimulation by region interaction [(*F*(6, 58190) = 12.20, *p* < .001)], such that accuracy is worse following cathodal stimulation to the PFC, during middle learning (*p* =.042) and late learning (*p* <.0001), when compared to sham stimulation (See Figure 4). Additionally, accuracy was better following anodal stimulation to the PFC during late learning (*p* = .002), compared to cathodal stimulation. Further, accuracy improved during the middle phase of learning following anodal stimulation to the cerebellum, when compared to cathodal stimulation (*p* = .043). No other effects reached significance (*F*s<2.01, *p*s>.16).

Next, we found an effect of phase [(*F*(3, 58190) = 165.19, *p* < .001)], though accuracy was best for early trials, follow by middle phase learning, and worse for late trials (*p* < .01). We should note accuracy was around ceiling (~90%) and this might be the result of fatigue/boredom, as RT clearly indicates participants were responding quicker in later learning phases. We also found a marginal effect of stimulation [(*F*(2, 148) = 3.00, *p* = .053)], such that accuracy was worse following cathodal stimulation, compared to sham (*p*= .050).

There was a significant phase by stimulation interaction [(*F*(6, 58190) = 23.15, *p* < .001)], such that accuracy was lower during middle (*p* = .006) and late (*p* < .001) learning following cathodal stimulation, compared to sham. However, accuracy was better during late (*p* =.008) learning following anodal stimulation, compared to cathodal. <u>Lastly</u>, a phase by region interaction [(*F*(3, 58190) = 11.95, *p* < .001)] emerged, in which accuracy was lower during the middle (*p* = .015) and late (*p* = .049) learning phases when stimulation was applied to the cerebellum, compared to the PFC. Taken together, accuracy is better following anodal stimulation to the PFC during later learning phases of learning, and anodal stimulation to the cerebellum during middle learning phases. However, performance during the middle and late phase of learning is worse following cathodal stimulation to the PFC.

### Balance

Means are displayed in Table 5 and visually depicted in Figure 5. We first examined sway area and sway deviation. We only found an effect of condition [(*F*(3, 446.17) = 30.64, *p* < .001)], such that sway area was larger in ECCB condition than ECOB (*p* < .001), EOCB (*p* < .001), and EOOB (*p*< .001) condition. No other effects of sway area reached significance (*F*s< 1.31, *p*s > .27). We next examined the sway deviation and found, for both SDX and SDY, sway deviation was larger for ECCB and EOCB, compared to ECOB (*p* < .001) and EOOB (*p* < .001). No other effects reached significance (*F*s < 1.88, *p*s > .083).

**Table 5.**
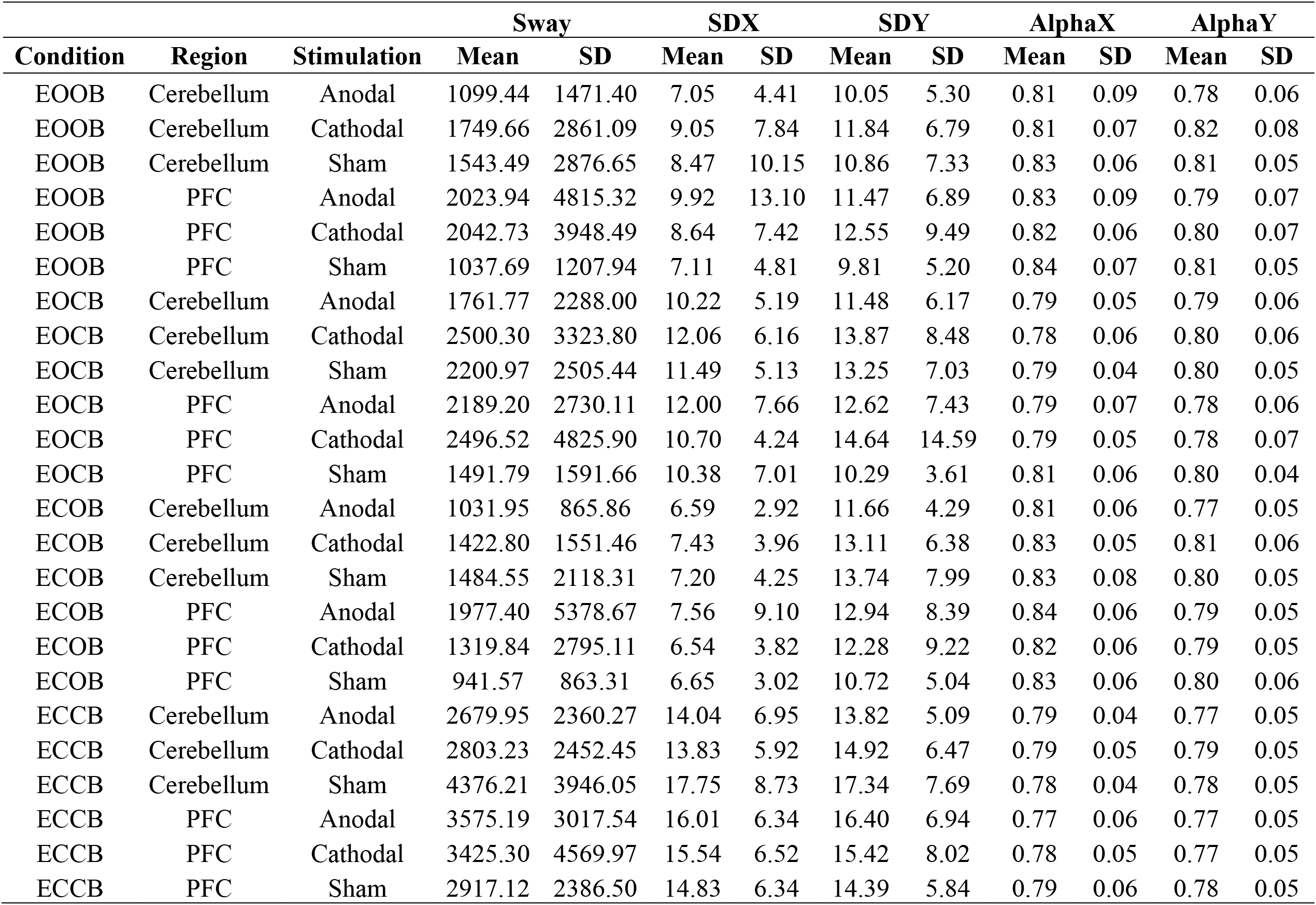
Mean complexity for the balance task by Condition, Region, and Stimulation Condition.

**Figure 5.**
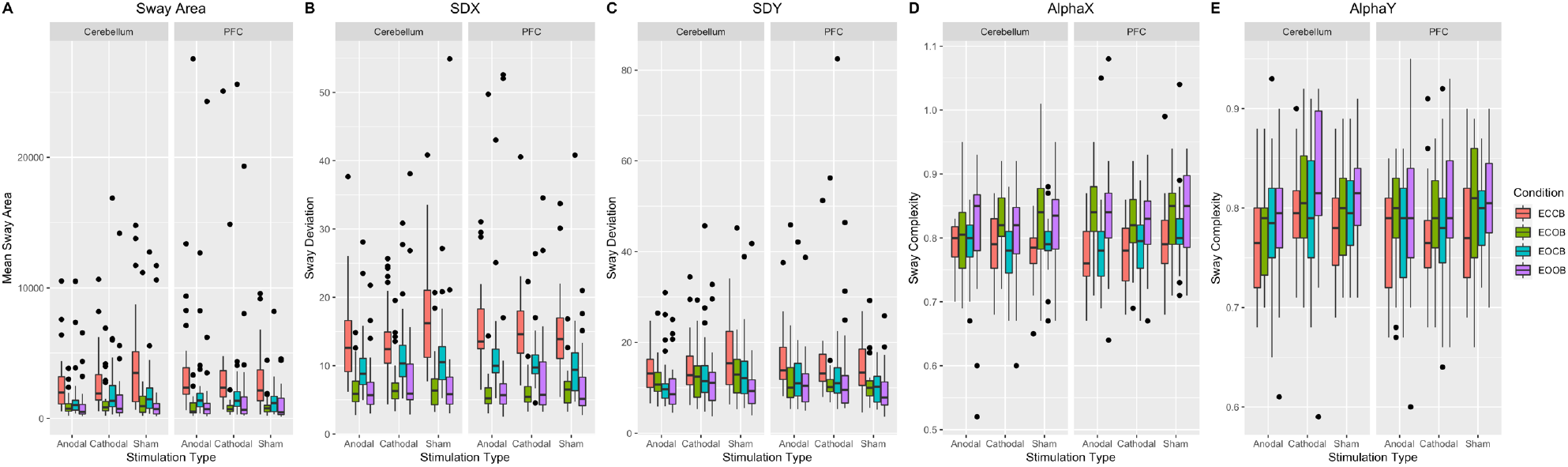
Mean complexity for the balance task by Condition, Region, and Stimulation Condition. Dots indicate outliers. Whiskers represent the interquartile range.

We next examined the complexity of the sway pattern. We found that AlphaX and AlphaY was lower for ECCB, compared to EOOB (*p* < .001) and EOCB (*p* <. 001). Taken together, we generally found larger sway area and deviation during conditions where eyes were closed (ECOB, ECCB), though complexity was lower during ECCB. However, we found no significant effects of stimulation or any interactions with stimulation.

## Discussion

The tDCS literature implicating the cerebellum in non-motor cognitive processing is growing, though past work using traditional tDCS over the right cerebellum to modulate cognitive performance has been mixed (Boehringer et al., 2013; Ferrucci et al., 2008; Majidi et al., 2017; Pope & Miall, 2012; Spielmann et al., 2017; van Wessel et al., 2016; Verhage et al., 2017), and few studies have taken advantage of HD-tDCS to examine the cerebellum in cognition (Ballard et al., 2019; Maldonado et al., 2019). There are several limitations to the current literature that may be contributing to the mixed results. First, traditional tDCS makes focal targeting more challenging. Second, both stimulation polarities have not been consistently used in the same studies, making comparison and interpretation across investigations more challenging. Third, it is an open question as to whether or not cognitive tasks are just more challenging to manipulate with tDCS, as effects on prefrontal stimulation are also somewhat mixed (Imburgio & Orr, 2018). To address these limitations, we used HD-tDCS to apply cathodal, anodal, or sham stimulation to either the PFC or the cerebellum, before participants completed motor (balance, sequence learning) and non-motor (Sternberg, Stroop) tasks. We predicted that cathodal stimulation would improve performance and anodal stimulation would impair performance, relative to sham, for both the cerebellum and PFC. Our findings suggest that anodal and cathodal stimulation both influence performance during working memory and sequence learning tasks, but not during the Stroop task or when testing postural sway, suggesting task specificity. These findings have broad implications for our understanding of cognitive processing in the cerebellum and how HD-tDCS might be used to further elucidate this understanding. These findings and their implications are discussed below.

### Sternberg Working Memory Performance

For verbal working memory performance, for both the cerebellum and PFC, we predicted increases in accuracy and decreases in reaction time following cathodal stimulation and predicted the opposite effect following anodal stimulation. We found that anodal cerebellar simulation negatively impacted reaction time. Interestingly, anodal stimulation improved accuracy under medium load, whereas cathodal stimulation hindered accuracy under high load. This does partially correspond with previous work by Ferrucci and colleagues (2008) who found both anodal and cathodal stimulation impaired performance on a Sternberg task. However, our work extends these findings to demonstrate a potential load effect. It is possible that a boundary exists where stimulation is beneficial, but this is dependent on stimulation type and task difficulty.

This further supports, and builds upon, the polarity specific nature of stimulation we demonstrate in sequence learning and its interaction with task difficulty. Specifically, when we examine accuracy, we demonstrate anodal stimulation improved accuracy under medium load, and cathodal stimulation hindered accuracy under high load. It is possible, that anodal stimulation might help cognitive processing during easier tasks, but not those that are automatic. Specifically, our work shows a benefit of anodal stimulation on medium load (three letters), but not low load (one letter). Most people are able to keep three items in memory with relative ease (Cowan, 2001; Miller, 1956); however, there is still a relative cognitive demand to maintain these items, while the one letter load allows for simple, and relatively automatic processing. Therefore, we suggest anodal stimulation helps maintain this level of cognitive demand. Under low load, one simply makes a dichotomous forced choice response that does not carry a demanding cognitive burden. In general, these findings provide further support for the cerebellar role in working memory, in conjunction with past work which strongly suggests stimulation to the cerebellum affects working memory ability (Ferrucci et al., 2008; Pope & Miall, 2012; Spielmann et al., 2017; Stoodley, 2012).

### Sequence Learning

For sequence learning, we predicted that cerebellar cathodal stimulation would improve learning, particularly during early learning, and anodal stimulation would hinder learning across all learning phases (Ballard et al., 2019). Cathodal stimulation slowed reaction times across learning phases, while anodal stimulation had no impact on reaction time. In regard to accuracy, we found anodal stimulation to both the PFC *and* the cerebellum improved accuracy, particularly in middle and late learning phases. However, cathodal stimulation to the PFC hindered accuracy in middle and late learning phases.

Unlike prior work showing performance improvements following cathodal stimulation (Ballard et al., 2019; Grimaldi et al., 2016), here we found cathodal stimulation had a negative impact on performance. For instance, when examining reaction time, we found that cathodal stimulation negatively affects reaction time, regardless of learning phase. Several have suggested distinct phases when learning new motor movements (Doyon, Gabitov, Vahdat, Lungu, & Boutin, 2018; Doyon et al., 1997; Karni et al., 1998). For instance, the cerebellum is particularly active in the early learning phase when procedural memories are created (Bernard & Seidler, 2013; Doyon et al., 2018), but cerebellar activations decrease as the cerebellum relies more on the newly created procedural models. It is possible that cathodal stimulation is disrupting the formation of these models, negatively impacting the execution of internal models downstream used to implement the previously learned procedures, negatively affecting reaction time. It is worth noting recent theories suggest motor related neural networks are used over the course of learning (Doyon et al., 2018). Critically, these networks, which include the cerebellum and cortical structures, are used to create internal models that are solidified by consolidation processes, which take time (Doyon et al., 2018; Doyon et al., 1997). Though speculative, it is possible cathodal stimulation, in this study, was hampering the processes needed for motor memory consolidation, ultimately resulting in poor accuracy in late learning phases (Doyon et al., 2018). However, the time window in the current study is short and more work would be needed to better understand whether memory consolidation was truly disrupted.

This polarity specific effect on learning phase might be crucial in future work, as our results suggest there might be a phase by stimulation interaction observed during sequence learning in the cerebellum. Previous work has suggested that working memory plays a significant role in motor learning (Bo & Seidler, 2009; Verwey, 1996; Verwey, 2001). It is suggested that the activation found in the cerebellum during motor learning tasks is working memory related, such that chunking might create internal models which are developed and implemented in order to learn a complete motor sequence (Bo & Seidler, 2009). Our results suggest cathodal stimulation might negatively impact the working memory process that are necessary during initial practice to learn novel motor sequences. Additionally, cathodal stimulation disrupts the automatic processes observed in late learning (Ramnani, 2014), and support a more automatic role for the cerebellum in later motor learning phases (Doyon et al., 2018; Karni et al., 1998).

### Stroop Task Performance

Contrary to our hypotheses, we did not find any effects of stimulation or region, only an effect of congruency when investigating inhibition using the Stroop task. Though limited, past work does suggest cerebellar tDCS might have an effect on inhibitory processes (Wynn et al., 2019). We suggest that relative simplicity of the current version of the Stroop task, where 50% of the trials were incongruent, might be contributing to this null finding. In the work conducted by Wynn and colleagues, approximately 11% of trails required an inhibitory response using a Go/No-go task. Though these tasks tap into similar underlying cognitive processes, the increased difficulty it the Go/No-go task used by Wynn and colleagues may require more input from the cerebellum, resulting in stimulation effect.

### Balance Performance

We next looked at the balanced data, where we predicted, for both the cerebellum and PFC, that cathodal stimulation would reduce sway area and variability and anodal stimulation would increase these variables (which would indicate poorer postural control). Consistent with the broader literature, we found larger sway area when eyes were closed, regardless of stimulation polarity or site (Bernard et al., 2014; Nardone et al., 1997; Nichols et al., 1995). However, no other effects emerged. Therefore, in the current study, it appears that stimulation over the right cerebellum or PFC does not have a major impact on balance performance in young adults. We should note that stimulation targeted the right cerebellum, as it is thought to be primarily involve in cognitive processes, and volume in this region has been shown to be positively related to balance performance in young and older adults (Bernard & Seidler, 2013).

Even though past work does suggest there is a relationship between cognition and postural control (Huxhold et al., 2006; Koziol et al., 2014), we did not directly stimulate the motor regions, such as the vermis, or more anterior regions of the cerebellum, perhaps resulting in this null effect. Even more, it is like that in healthy young adults, balance performance is optimal, which leaves little room for improvement and any perturbation may not be enough to create performance decrements. However, future work may continue to investigate whether there is a direct relationship between motor and nonmotor regions of the cerebellum when it comes to balance, particularly in those who might be greater fall risks, such as older adults.

### Limitations

HD-tDCS is a relatively new methodology whose main purpose is to better direct the current being applied to the scalp. As such, there has been little work using this approach to investigate behavior, to this point, particularly with respect to the cerebellum. Even with improved targeting, it is possible that the stimulation montage might affect regions outside of the desired target area, impacting behavior in an unpredicted way.

As noted above, cognitive task difficulty may be impacting our findings here. Even though 50% of the trials in the Stroop task were incongruent, participants might have mastered the task quickly enough such that the majority of the time they spent completing the task was on “autopilot”. Thus, the PFC processed the information with no need to recruit additional resources. It is possible that the cerebellum did not play a major role in processing information to complete this task as the task became too simple, as there was no real cognitive demand. Additionally, the high load on the Sternberg tasks only required participants to remember 5 letters, which might not be challenging enough to get a good deal of performance variability. Future work with more complex tasks is of great interest, as past work has demonstrated that the cerebellum is involved in inhibition (Schmahmann & Sherman, 1998; Wynn et al., 2019) and verbal working memory (Ferrucci et al., 2008; King et al., 2019; Pope & Miall, 2012; Stoodley, 2012).

## Conclusion

The current study applied HD-tDCS to both the cerebellum and PFC to better understand the relative necessity of the cerebellum in non-motor cognitive processing. Our findings suggest support a role for the cerebellum in verbal working memory, but not inhibition. However, the latter may be due to task difficulty. Furthermore, stimulation might be load specific when applied to the cerebellum, such that the effect of stimulation might differ depending on the cognitive demands of the task. We found anodal stimulation negatively impacts reaction time during effortful processing, but is beneficial for accuracy on less effortful processing. Further, cathodal stimulation negatively effects processing in both the cerebellum and PFC. This provides initial evidence to suggest the cerebellum is involved in non-motor cognitive processing, but influence might be dependent on task difficulty. Future work should continue to explore the interaction between cerebellar involvement and cognitive demand to further elucidate the necessity of the cerebellum and non-motor cognitive processing.

## Acknowledgements

We would like to thank research assistants Cassidy Carrasco, Malin Chambers, Sydney Eakin, Kristin Eiland, James Goen, Abigail Miller, Isai Ramirez, Lisset Salinas for their help with data collection.

## Declarations

This research did not receive any specific grant from funding agencies in the public, commercial, or not-for-profit sectors. The authors have no relevant financial or non-financial interests to disclose. The authors do not have any conflicts of interest.

